# Degradation factor 1, Def1, regulates mRNA translation and decay through Ccr4-Not-dependent ubiquitylation of the ribosome

**DOI:** 10.64898/2025.12.10.693450

**Authors:** O.T. Akinniyi, A. Sebastian, S Kulkarni, I Albert, J.C. Reese

## Abstract

Yeast Def1 is well known for its role in regulating RNA polymerase II elongation and degrading the large subunit of polymerase during transcriptional stress. It is an abundant cytoplasmic protein that undergoes stress-induced processing and is then transported to the nucleus. Previous research from our lab has shown that Def1 interacts with various proteins involved in mRNA decay and translation control, and that it regulates mRNA half-lives, suggesting an important role in the cytoplasm. In this study, we report that Def1 binds polyribosomes and that its null mutant strain exhibits phenotypes indicating a role in translation. Ribo-seq analysis revealed that deleting *DEF1* altered ribosome footprints on mRNAs and increased the dwell time of ribosomes at non-optimal codons in the A-site. Additionally, results from a codon-optimality reporter assay suggest that Def1 facilitates the degradation of mRNAs containing non-optimal codons. The Ccr4-Not complex links codon optimality to mRNA decay, and Def1’s binding to ribosomes depends on its ubiquitin-binding domain, as well as the ubiquitylation of eS7a in the small ribosomal subunit by the Ccr4-Not complex. Moreover, the polyglutamine-rich, unstructured C-terminus of Def1 is crucial for its interaction with RNA decay and translation factors. This indicates that Def1 functions as a ubiquitin-dependent scaffold, connecting translation status to mRNA decay. In summary, we have identified a cytoplasmic function for Def1 in translation and established it as a regulator of gene expression, spanning both transcription and translation processes.

## Introduction

Cells regulate multiple steps in gene regulation when they encounter stress, such as DNA damage. This includes processes from transcription to translation, during which nucleic acid damage interrupts gene expression, allowing repair and enabling the cell to resume normal functions. During transcription, damage to the DNA template halts RNA polymerase II (RNAPII) progression. If the damage cannot be repaired, RNAPII is targeted for destruction through ubiquitylation and degradation of its largest subunit, Rpb1. (Wilson et al. 2013a; Steurer and Marteijn 2017; Akinniyi and Reese 2021; Reese 2023).

Several factors contribute to Rpb1 degradation, including yeast Degradation Factor 1 (DEF1), a key player in this process. Def1 facilitates the sequential recruitment of ubiquitin ligases and other proteins necessary for the degradation of RNA polymerase II (RNAPII).(Woudstra et al. 2002; Wilson et al. 2013b; Akinniyi and Reese 2021). This function depends on its ubiquitin-binding domain (Wilson et al. 2013b). Def1 is believed to have broader functions in transcription, including regulating TFIIH and participating in transcription initiation and elongation (Somesh et al. 2005; Damodaren et al. 2017; Akinniyi and Reese 2021). Paradoxically, Def1 is a highly abundant cytoplasmic protein. It was later discovered that a portion of Def1 undergoes DNA damage-dependent proteasomal processing in the cytoplasm, separating the N-terminus of the protein from the unstructured, polyglutamine-rich C-terminus. (Wilson et al. 2013b). The C-terminus of Def1 keeps it in the cytoplasm. It remains uncertain whether Def1 has functions in the cytoplasm or if its presence there simply prevents interference with RNAPII transcription in the nucleus.

Recently, we showed that deleting *DEF1* repressed mRNA synthesis, consistent with its established role in transcription. However, we also observed a global reduction in mRNA degradation rates (Akinniyi et al. 2025). Additionally, proximity labeling experiments using BioID indicated that Def1 predominantly tagged cytoplasmic proteins, especially those involved in mRNA decay and translation, including the Ccr4-Not deadenylase complex. (Pfannenstein et al. 2024; Akinniyi et al. 2025; Kulkarni et al. 2025). A different BioID study found Def1 located near the small subunit of the ribosome. (Schmitt and Valerius 2019; Schmitt et al. 2021). Moreover, the human homolog of Def1, UBPA2/2L, binds various cytoplasmic RNA-binding proteins, including the human Ccr4-Not complex, and interacts with ribosomes (Youn et al. 2018; Luo et al. 2020; Kulkarni et al. 2025). These observations indicate that Def1 plays a role in the posttranscriptional regulation of mRNAs in the cytoplasm, including in translation.

Ccr4-Not was first identified as a transcriptional regulator in yeast, but it is now known to function at many points of gene control, including transcription, mRNA deadenylation, translation quality control, and protein ubiquitylation(Kruk et al. 2011; Miller and Reese 2012; Collart 2016; Chalabi Hagkarim and Grand 2020). Ccr4-Not is the primary mRNA deadenylase responsible for initiating degradation by removing the poly(A) tail. It also promotes transcription elongation and plays a crucial role in the DNA damage-dependent degradation of Rpb1, the large subunit of RNA polymerase II, a function that it shares with Def1 (Kruk et al. 2011; Reese 2013; Jiang et al. 2019). The Not4 subunit of Ccr4-Not functions as a RING domain ubiquitin ligase, although the full range of substrates modified by Not4 is not yet known. One of the first proteins identified as ubiquitylated by Not4 is eS7a (Rsp7a), a ribosomal small-subunit protein (Panasenko and Collart 2012; Buschauer et al. 2020; Allen et al. 2021; Müller et al. 2025). Ccr4-Not has been found to play an important role in the co-translational degradation of poorly translated mRNAs, such as those that contain a high proportion of non-optimal codons. When a ribosome stalls over a non-optimal codon in its A-site, the tRNA in the E-site is evicted. This site is then occupied by the N-terminus of the Not5 subunit of the Ccr4-Not complex. Following this, Not4 ubiquitylates eS7a, and the deadenylase activity of Ccr4-Not degrades the mRNA transcript.(Buschauer et al. 2020; Collart et al. 2023; Müller et al. 2025). While the Not4-dependent ubiquitylation of the small subunit is linked to translation quality control and co-translational decay, the identity of a factor that recognizes this modification and triggers these downstream events remains unknown.

Here, we find that Def1 associates with polyribosomes, a process that depends on its ubiquitin-binding domain and on Not4-dependent ubiquitylation of the small ribosomal subunit. We present evidence that the C-terminus of Def1 recruits translation control and mRNA decay factors to the ribosome, which helps regulate mRNA decay based on codon optimality. Therefore, we have identified a novel function for Def1 and revealed a previously unknown reader of ribosomal protein ubiquitylation.

## Results

### Def1 associates with polyribosomes

Our previous BioID studies showed that Def1-TID labeled proteins include those involved in translation and mRNA decay (Akinniyi et al. 2025). Translation and RNA decay are coupled processes (Hu et al. 2009; Bae and Coller 2022; Wu and Bazzini 2023); therefore, we hypothesized that Def1 functions in translation and/or interacts with polyribosomes. To investigate this, we conducted polysome profiling to determine whether Def1 associates with polyribosomes. We fractionated cell extracts using a 10-50% sucrose gradient and probed the fractions with anti-Def1 antibody. Our findings revealed that the majority of Def1 co-migrated with the mono- and polyribosome fractions, while only a small amount was found in the mRNP fraction (Fig. 1A). Def1 was also detected in the fractions containing the 40S and 60S ribosomal subunits. To verify that Def1 is associated with polysomes, we prepared lysates and gradients in buffers containing EDTA, a known polysome-disassembling agent. Treatment with EDTA disrupted Def1 migration in the heavier polysome fractions (Fig. 1B), providing evidence that Def1 associates with polyribosomes. Interestingly, Def1 still co-migrated with the 40S and 60S ribosomal subunits, indicating that it interacts with the small and large ribosomal subunits independently of translation.

**Figure 1:**
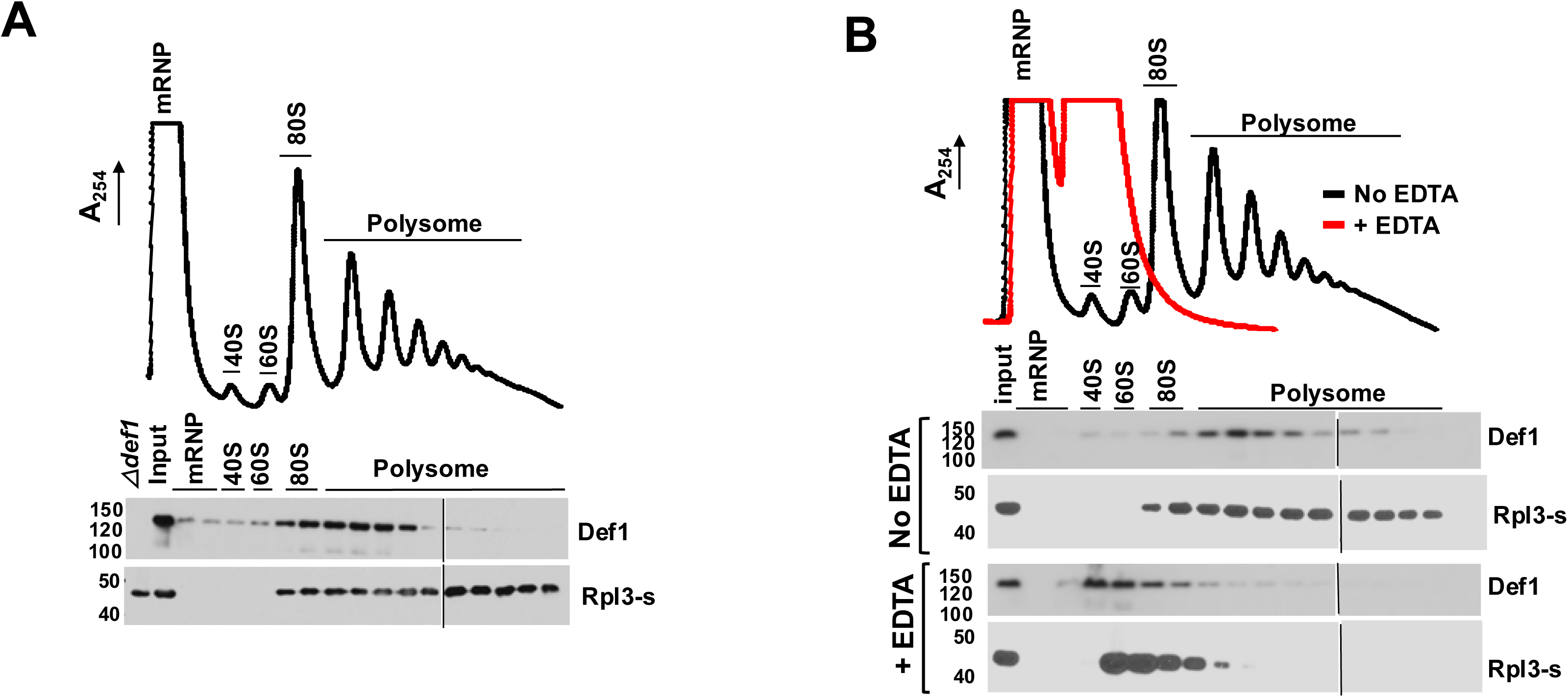
Def1 associates with polyribosomes. (A). Polysome profiling. Cell lysate was fractionated on a 10-50% sucrose gradient as described in the Methods section. The A_254_ recording of the gradient fractions is displayed on the y-axis. Def1 and Rpl3-s (large subunit protein) were detected by Western blotting. The locations of the mRNP, 40S, 60S, 80S monosomes and polysomes are indicated. Two blots were used to analyze all the fractions, and the splice is indicated with a line. An extract from a *def1Δ* strain was analyzed to validate the antibody. (B) Migration of Def1 in the presence (red) or absence (black) of the polysome-disrupting agent EDTA. EDTA was added to the extract and gradient to disrupt polysomes.

The association of a protein with ribosomes implies that it influences translation. To investigate the impact of deleting the *DEF1* gene on translation, we first assessed the sensitivity of the *def1Δ* mutant to hygromycin B, a translation-inhibiting drug. Indeed, the mutant was sensitive (Fig. 2A). Puromycin incorporation into polypeptides is a standard method for measuring translation (Cary et al. 2014; Enam et al. 2020). Consequently, we evaluated protein synthesis by examining puromycin incorporation in both wild-type and *def1Δ* strains. Our results showed a significant reduction of puromycin incorporation, approximately 2.5-fold, in the mutant compared to the wild-type cells (Fig. 2B).

**Figure 2.**
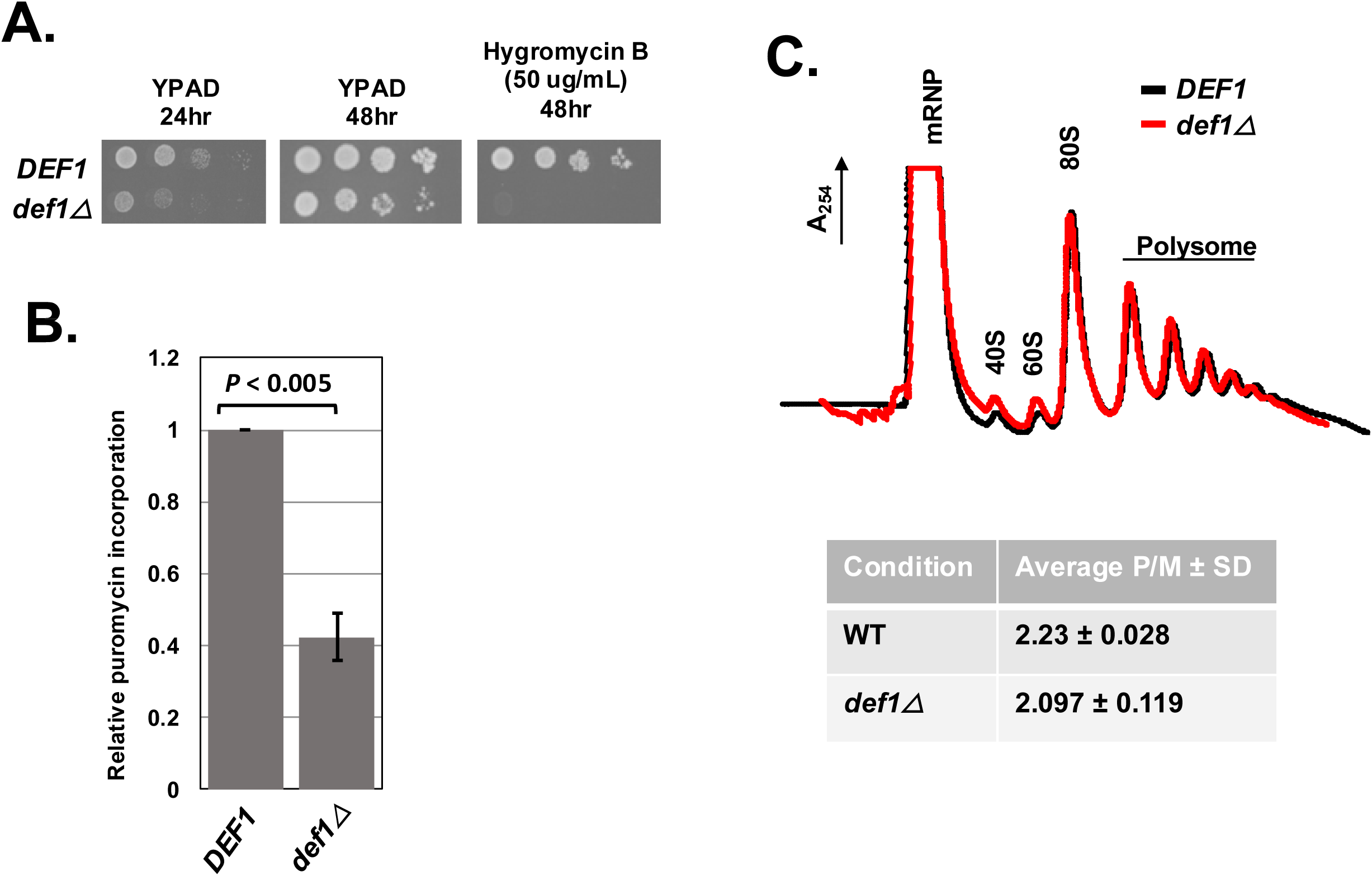
Def1 is required for translation. (A) *DEF1* null cells are sensitive to translation inhibitor, hygromycin B. Cells were spotted onto YPAD media, and the same with 50 ug/ml hygromycin. Images were captured at 24- and 48-hr. (B) Puromycin incorporation is impaired in *def1Δ*. Cells were grown to log phase, and puromycin was added for 15 min before harvesting. Puromycin incorporation was detected by Western blot analysis of cell extracts using anti-puromycin antibody. Results are the average and standard deviation of biological triplicates (N=3). (C). Representative polysome profiles of wild-type and *def1Δ* cells overlayed (wild-type cells in black and *def1Δ* in red). The average polysome-to-monosome ratios (P/M) were determined and represented in the table. The area under the curve (AUC) for the monosome (80S) and polysome fractions was calculated using PeakChart 3 software (Brandel). The average and standard deviations (N=3) of independent experiments are shown.

Mutations in certain translation factors lead to altered polysome-to-monosome ratios (P/M) in polysome profiles. However, when we compared the polysome profiles of wild-type and *def1Δ* cells, we found no detectable differences in the peaks of 40S, 60S, and 80S ribosomal subunits, nor in the polysome fractions (Fig. 2C). Therefore, although the mutant cells did not show any change in the polysome-to-monosome ratio, the association of Def1 with polysomes, along with the mutant’s sensitivity to translation inhibitors and decreased puromycin incorporation, suggests that Def1 plays a role in translation.

### Ribosome sequencing (Ribo-seq) reveals a function of Def1 in translation

Bulk polysome profiling can obscure changes in translation status, including transcript-specific regulation, initiation-elongation dynamics, and ribosome-dwelling kinetics. We therefore used ribosome sequencing (Ribo-seq) to investigate translation changes in *Δdef1* cells. We mapped over 12 million ribosome-protected footprints (RPFs) and mRNA reads for wild-type cells and over 19 million for *def1Δ* cells, totaling 5,094 transcripts (Supplemental file 1). The Ribo-seq data were analyzed using the RiboMiner toolset (Li et al. 2020). The three replicates for each experimental condition showed strong correlation and clustered together in PCA plots and the RPFs exhibited strong three-nucleotide periodicity, a characteristic of successful Ribo-seq execution (Supplemental Fig. S1A, B).

Metaplots of ribosome distribution across mRNAs were created and centered over the start and stop codons (Fig. 3A and B, respectively). Ribosome density peaked at both the start and stop codons in both cell types, which was expected. However, ribosome protected fragments (RPFs) showed a significant reduction between codons 30-45, along with a slight increase in density observed over the start codon in the *def1Δ* mutant (Fig. 3A). This same pattern persisted when the metaplots were normalized to open reading frame (ORF) length, suggesting that the reduction is not due to changes in density over shorter ORFs (refer to Supplemental Fig. S1C). There were no noticeable differences in the RPFs near the stop codon. Another way to analyze shifts in ribosome density across mRNAs is through a polarity plot, where negative and positive scores indicate a bias towards the 5’ and 3’ ends, respectively (Vorontsov et al. 2021). We observed that the *def1Δ* mutant exhibited a shift in the density of ribosome footprints toward the 3’ end compared to wild type cells, resulting in a more positive value (Fig. 3C). According to the metaplot analysis, this shift seems to be driven primarily by a reduction in ribosome footprint (RPF) density in the first 30-45 codons, rather than an increase further along the transcript. This decrease in density occurs at codons corresponding to the amino acids expected to be located within the ribosome’s peptide exit channel (Wilson et al. 2016; Dao Duc and Song 2018; Gamerdinger et al. 2019). This complex pattern may be explained by Def1p’s role in stabilizing ribosomes as the polypeptide moves through the exit channel, possibly by preventing premature translation termination. Additionally, we observed a recovery in ribosome density within the transcript and a slight increase in density at the 3’ end in the mutant, which may result from slowed ribosomal elongation. This slowed elongation could account for the reduced puromycin incorporation observed in the mutant (Fig. 2B).

**Figure 3.**
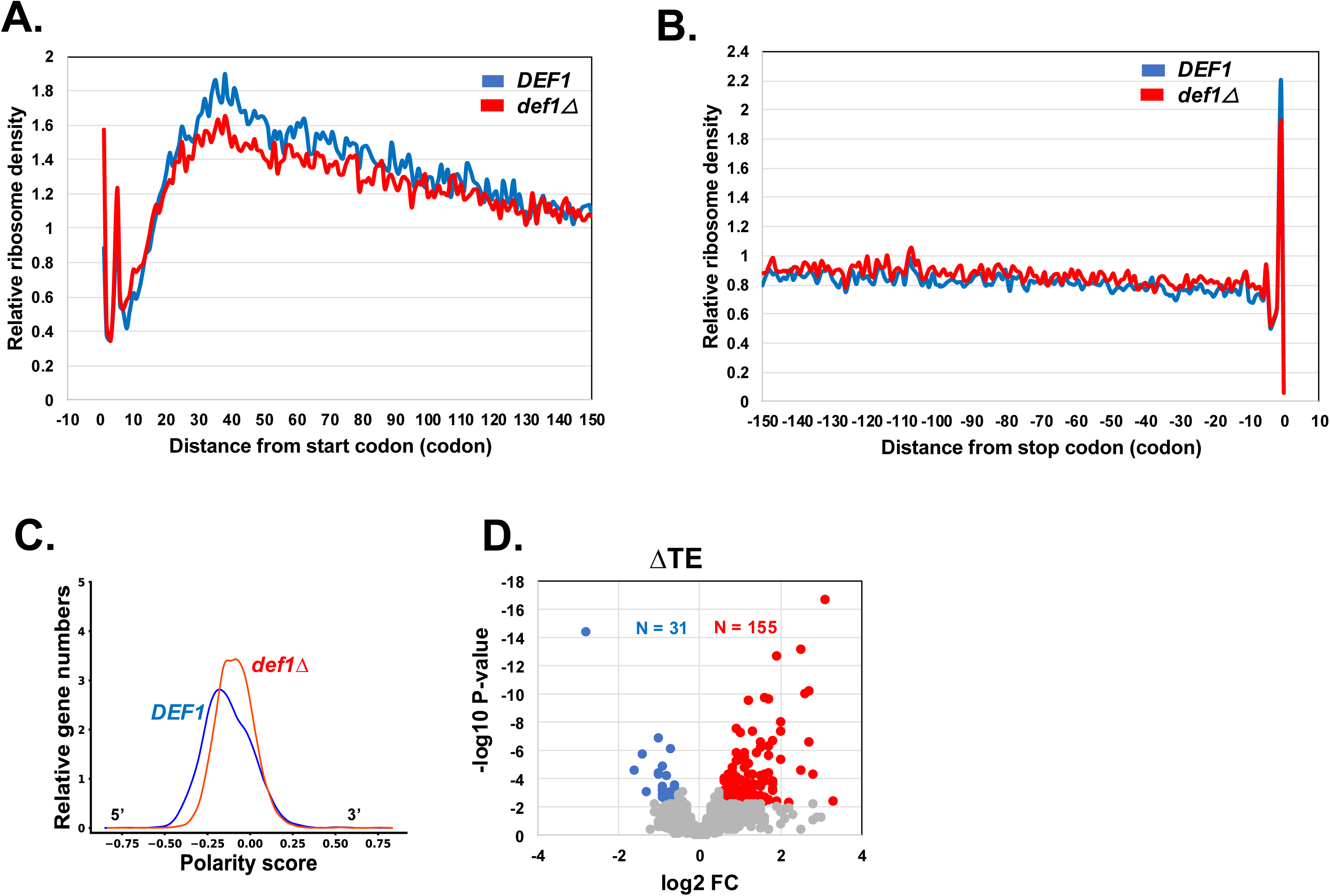
Ribosome profiling (Ribo-seq) of the *def1Δ* mutant revealed altered ribosome distribution and increased non-optimal codons in the A-site. (A) Metaplot analysis of ribosome density distribution over the first 150 codons of all 5140 transcripts. Metagene plots were generated using RiboMiner v0.2 and the ribosome density at each position is normalized by the overall ribosome density within the coding sequence. Wild type (in blue) and *def1Δ* (in red) conditions are plotted. (B) Metaplot analysis as in *A*. The ribosome density distribution across 150 codons upstream of the stop codon of all ORFs. (C) Polarity score of ribosome density across mRNAs. Transcriptome-wide shift in overall ribosome distribution was evaluated and represented for both wild type (in blue) and *def1Δ* mutant (in red). (D) Change in translation efficiency (TE) in a *def1Δ* mutant.

We next examined changes in translation efficiency (TE), defined as ribosome density per mRNA, adjusted for mRNA levels. TE can help identify transcripts that are particularly sensitive to Def1 loss. However, an increase in TE for certain mRNAs can result from either increased translation efficiency or ribosome stalling, since ribosome profiling (Ribo-seq) measures ribosome density rather than directly measuring translation activity. Of the 5,094 mRNAs detected, 155 transcripts showed an increase in TE, while 31 transcripts demonstrated a decrease when applying a cutoff of greater than 1.5-fold change (FC) and a p-value of less than 0.005 (Fig. 3D). Gene Ontology (GO) term analysis of the 155 mRNAs revealed enrichment in genes that regulate macromolecule metabolism pathways (Supplemental Fig. S1D). In contrast, no specific GO terms were associated with the mRNAs that exhibited reduced TE.

### Codon-dependent regulation of translation and decay

We investigated the impact of Def1 on ribosome dwell time at specific codons. Certain factors that interact with Def1, such as Ccr4-Not, are known to regulate ribosome arrest at non-optimal codons (Buschauer et al. 2020; Collart et al. 2023; Müller et al. 2025). We categorized codons based on their codon stabilization coefficient (CSC) into three equal groups: high, intermediate, and low codon optimality. Then, we calculated the relative occupancy of codons in the A-site of ribosomes in mutant cells compared to wild-type cells. Notably, we observed a significant increase in A-site occupancy for the intermediate and non-optimal codons in the mutant cells when compared to the optimal codon group (Fig. 4A). When we plotted the changes in ribosome density at specific codons, we found a general mild increase in density across most codons. However, consistent with the box plots in Fig. 4A, the majority of non-optimal codons showed enhanced A-site occupancy in the mutant (Supplemental Fig. S1E). These findings suggest that Def1 improves the ribosome’s capacity to translate less optimal codons.

**Figure 4.**
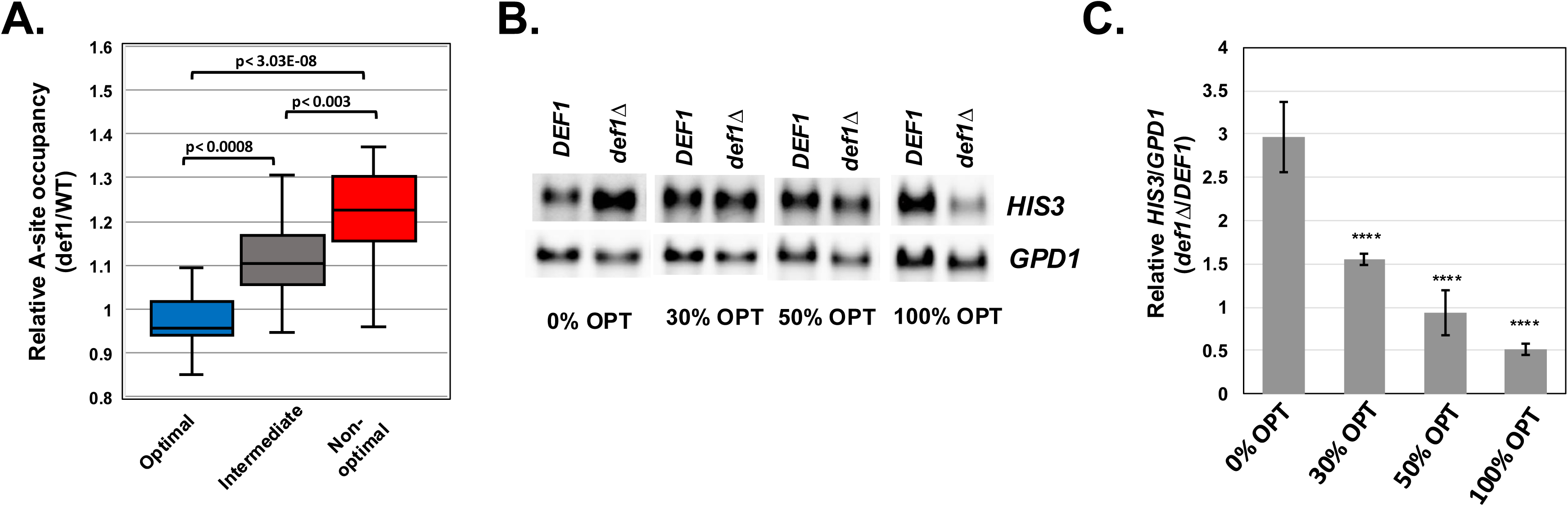
Def1 regulates codon-dependent mRNA decay. (A). Change in ribosome density over the A-site versus codon optimality. Codons were separated into three bins based on CSC scores: optimal (n=20), intermediate (n=21), and non-optimal (n=20). Stop codons were not included in this analysis. The change in ribosome density over the A-site obtained in the Ribo-seq experiment is plotted on a log2 scale. P-values were calculated using a Mann-Whitney test. (B). Representative Northern blot of codon optimality reporter gene assay. *HIS3* ORFs with variable codon optimality (indicated below each panel) were placed under the control of the *GPD1* promoter. The endogenous *GPD1* mRNA was used as a loading control and to correct for changes in promoter activity in the mutant. The image capture times for each *HIS3* derivative varied because of differences in steady state levels: 150s for 0% and 30% OPT, 60s for 50% OPT, and 20s for 100% OPT (C). Averages and standard deviations of reporter mRNA expression. The signals for the *HIS3* mRNA were first normalized to the endogenous *GPD1* mRNA, and then the values in the *def1Δ* cells were divided by the values in *DEF1* cells (relative normalized signal) (n=4). P-values were calculated using a two-tailed, unpaired t-test, and the values of the 0% reporter were compared to those of the other constructs. ****, p<0.0001.

mRNAs that are enriched in non-optimal codons exhibit decreased stability through a process known as co-translational decay (Roy and Jacobson 2013; Inada and Beckmann 2024; Müller et al. 2025). This process has been assessed by analyzing the half-lives of *HIS3* variants with varying codon optimality expressed from the *GAL1* promoter after transcription shutdown (Radhakrishnan et al. 2016). We attempted to use these reporter constructs for our shutdown experiments; however, the *def1Δ* mutant showed defective expression from the *GAL1* promoter (Akinniyi, O.T. and Reese, J.C, unpublished). Therefore, we replaced the *GAL1* promoter with the constitutive *GPD1* promoter. Given that deleting *DEF1* reduces mRNA transcription (Akinniyi et al. 2025), we normalized the expression of the *GPD1* promoter-driven *HIS3* reporter to the chromosomal *GPD1* gene to correct for changes in promoter activity. The values from the mutant cells were then normalized to those from wild-type cells. Since all reporter constructs are controlled by the same promoter, the relative change in the steady-state *HIS3* mRNA level reports on mRNA stability.

To validate this new reporter system, we analyzed a *not4ringΔ* mutant, which is defective in codon-dependent co-translational mRNA decay (Buschauer et al. 2020). As expected, the relative amount of the 0% optimal reporter in the *not4ringΔ* mutant was higher compared to the 50% and 100% optimal constructs (see Supplemental Fig. S2). In other words, the unstable 0% optimal reporter was expressed better; and thus, had greater stability in the mutant compared to wild-type cells. This can also be observed in Northern blots (Supplemental Fig S2). Having confirmed the reporter assay with a known co-translational decay mutant, we proceeded to analyze the *def1Δ* mutant. Similar to the *not4ringΔ* mutant, the relative steady-state levels of the reporter decreased as codon optimality increased (Fig. 4B and C). There was approximately a five-fold difference in the relative expression between the 0% and 100% optimal reporters in the mutant. This finding indicates that non-optimal mRNAs are more stable in mutant cells compared to wild-type cells. Thus, our reporter assay suggests that Def1 is involved in codon-dependent co-translational decay.

### Def1 binding to polysomes requires its ubiquitin-binding domain

We investigated which region of Def1 is necessary for ribosome binding. Def1 has an N-terminal CUE domain that interacts with ubiquitin-homology domains and ubiquitin, as well as a C-terminal polyglutamine-rich (polyQ) domain that is essential for its accumulation in the cytoplasm (Fig. 5A) (Wilson et al. 2013b). To study this, we generated a strain expressing a truncation mutant of Def1 (Def1_1-380_) that lacks the polyQ region and contains GFP fused to the C-terminus. We also generated and analyzed full-length Def1-GFP (1-738) as a positive control.

**Figure 5:**
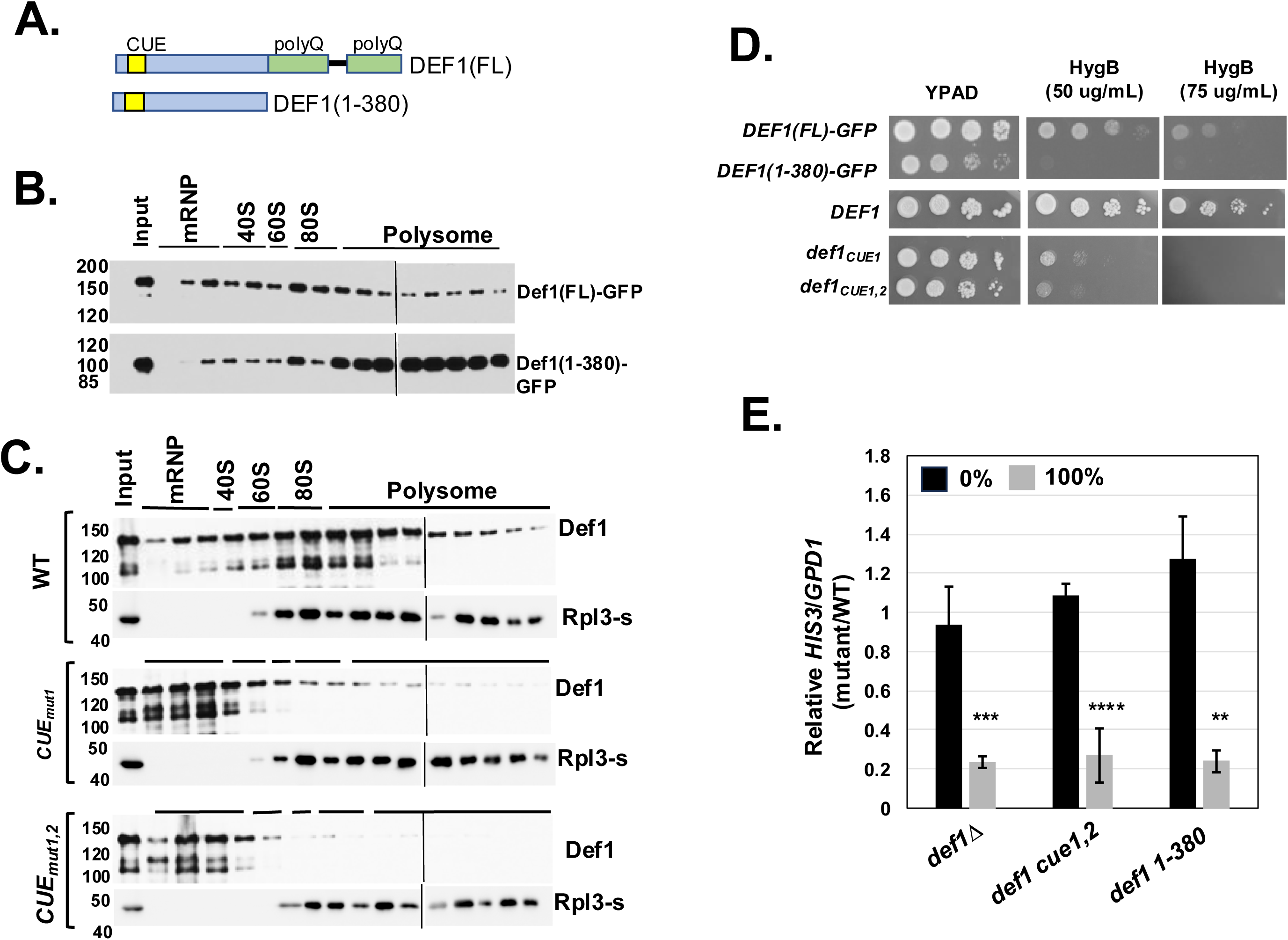
The ubiquitin-binding domain (CUE) of Def1 is required for ribosome binding. (A). Schematic of the Def1 protein with the CUE and polyQ domains illustrated. (B). Def1(1-380)-GFP migration in polysome profiles. Western blotting was performed using anti-GFP antibodies. (C). Polysome fractions of wild type, CUE_mut1,_ and CUE_mut1,2_ versions of Def1. The amino acid substitutions are described in Supplemental Table 1 and (Wilson et al. 2013b). Western blotting was performed using antiserum to Def1 and Rpl3-s. (D) Hygromycin B (HygB) sensitivity growth assay. As in Fig. 2A, except that two concentrations of drug were used. (E). *HIS3* codon optimality assay. The assay was conducted as described in Figure 4C. 0% (black bars) and 100% (gray bars) refer to the codon optimality of the *HIS3* construct. Biological triplicates were analyzed (n=3). **, p<0.01; ***, p<0.001; ****, p<0.0001.

Our findings showed that the Def1 (1-380) mutant retained ribosome binding, and moreover, we reproducibly observed enhanced polysome binding, especially in the heavier (larger) polysome fraction (Fig. 5B). One possible explanation for the increased polysome interaction in the polyQ truncation mutant is that bioinformatics tools predict a significant intrinsically disordered segment within the polyQ region (Akinniyi, O.T. and Reese, J.C, unpublished). Additionally, Def1 has been shown to form polyQ-dependent amyloid-like filaments *in vitro* (Damodaren et al. 2017). It is possible that a portion of full-length Def1 is trapped in an inaccessible cellular fraction, and removing the polyQ domain may increase the amount of Def1 available for ribosome binding. Another possibility is that the polyQ region recruits factors that influence translation elongation (see below). Nevertheless, our results indicate that the Q-rich C-terminus of Def1 is not required for its association with ribosomes.

Since Def1(1-380) binds to ribosomes, we hypothesized that the ubiquitin-binding CUE domain is responsible for this interaction. To test our hypothesis, we employed CRISPR-Cas9 gene editing to introduce two clusters of mutations in conserved residues within the CUE domain, labeled CUEmut1 and CUEmut2. These mutations disrupt Def1’s binding to ubiquitin and ubiquitin-homology domains (Wilson et al. 2013b). Using strains that express these mutant versions of Def1 (CUE_mut1_ and the CUE_mut1,2_ double mutant), we observed that disrupting the ubiquitin binding function of the CUE domain significantly reduced Def1’s association with ribosomes (Fig. 5C). Interestingly, polysome profiles shown in Figure 1B indicated that treatment with EDTA disrupted Def1’s association with polysomes, but not with the 80S, 60S, or 40S ribosomal fractions, with minimal Def1 present in the mRNP fractions. In contrast, mutations in the CUE domain led to a shift of most Def1 protein into the mRNP fraction. These results suggest that the CUE domain and its ubiquitin-binding function are essential for Def1’s association with polyribosomes and individual ribosomal subunits.

We screened the Def1 mutants for their sensitivity to the translation elongation inhibitor hygromycin B. The strain expressing Def1(1-380) exhibited sensitivity to hygromycin B similar to the *def1Δ* strain (Fig. 5D). Additionally, the CUE mutant strains also showed sensitivity to the translation inhibitor, but to a lesser extent than the null mutant; the CUE mutants required a higher concentration of the drug to inhibit growth compared to the deletion mutant (Fig. 5D vs. 2A).

We next determined whether the CUE domain and the C-terminal polyQ region of Def1 are required for co-translational decay using the codon-optimality reporter assay. The results shown in Figure 5E indicate that both the CUEmut_1,2_ and Def1 (1-380) mutants were defective for co-translational decay (Fig. 5E). As another test, we used an MS2/MCP tethering reporter assay. We previously demonstrated that tethering wild-type Def1 to a reporter mRNA decreased reporter expression (Akinniyi et al. 2025). In the current study, we expressed the CUE and Def1 (1-380) mutants fused to MCP and compared their effect on reporter expression to that of the full-length Def1 fused to MCP. Our findings show that all mutants exhibited impaired suppression of the reporter mRNA (Supplemental Fig. S3). Overall, these results demonstrate that both the ubiquitin-binding CUE domain and the polyQ-rich segment of Def1 are essential for the post-transcriptional regulation of mRNAs.

### Not4-mediated ubiquitylation of eS7a (Rps7A) is required for the association of Def1 with polyribosomes

Ubiquitylation of ribosomal proteins (RPs) has several important roles in translation, including rescuing stalled ribosomes and degrading faulty nascent polypeptides through the ribosome quality control (RQC) pathway (Inada and Beckmann 2024). Based on our findings that the CUE domain of Def1 is essential for its association with polysomes, we hypothesized that the ubiquitylation of an RP contributes to its recruitment. Multiple ubiquitin ligases modify RPs, including Not4, a subunit of the Ccr4-Not complex (Panasenko and Collart 2012; Ikeuchi et al. 2019; Allen et al. 2021; Inada and Beckmann 2024). As previously mentioned, BioID experiments suggested that Def1 and the Ccr4-Not complex are located in close proximity, and that Def1 and Not4 exhibit both genetic and physical interactions (Jiang et al. 2019; Pfannenstein et al. 2024). Not4 ubiquitylates eS7A (Rps7A) on arrested ribosomes, particularly those stalled due to non-optimal codons in the A-site (Buschauer et al. 2020; Inada and Beckmann 2024; Müller et al. 2025).

First, we investigated whether Ccr4-Not is necessary for Def1 binding to ribosomes by depleting Not1, the scaffold subunit of the complex. Since Not1 is an essential gene, we employed an auxin-inducible degron strategy to deplete it from cells (Shetty et al. 2019). In the absence of auxin, the Not1-AID protein was expressed alongside the native protein; however, the addition of auxin led to its depletion (Fig. 6A). We found that depleting Not1 resulted in a significant decrease in the association of Def1 with ribosomes, as well as its shift into the mRNP fraction (Fig. 6B). Additionally, either deleting the *NOT4* coding sequence or depleting the Not4 protein through auxin-dependent degradation disrupted Def1’s association with polyribosomes (Supplemental Fig. S4). Next, we determined whether the ubiquitin ligase activity of Not4 is required for Def1 recruitment to ribosomes. We analyzed a Not4 RING domain deletion mutant (*not4ringΔ*), which lacks ubiquitin ligase activity but retains the overall integrity of the Ccr4-Not complex (Jiang et al. 2019). Here, too, we observed a significant reduction in Def1 binding to ribosomes, indicating that Not4’s ubiquitin ligase activity is indeed required for Def1 binding to ribosomes (Fig. 6C).

**Figure 6.**
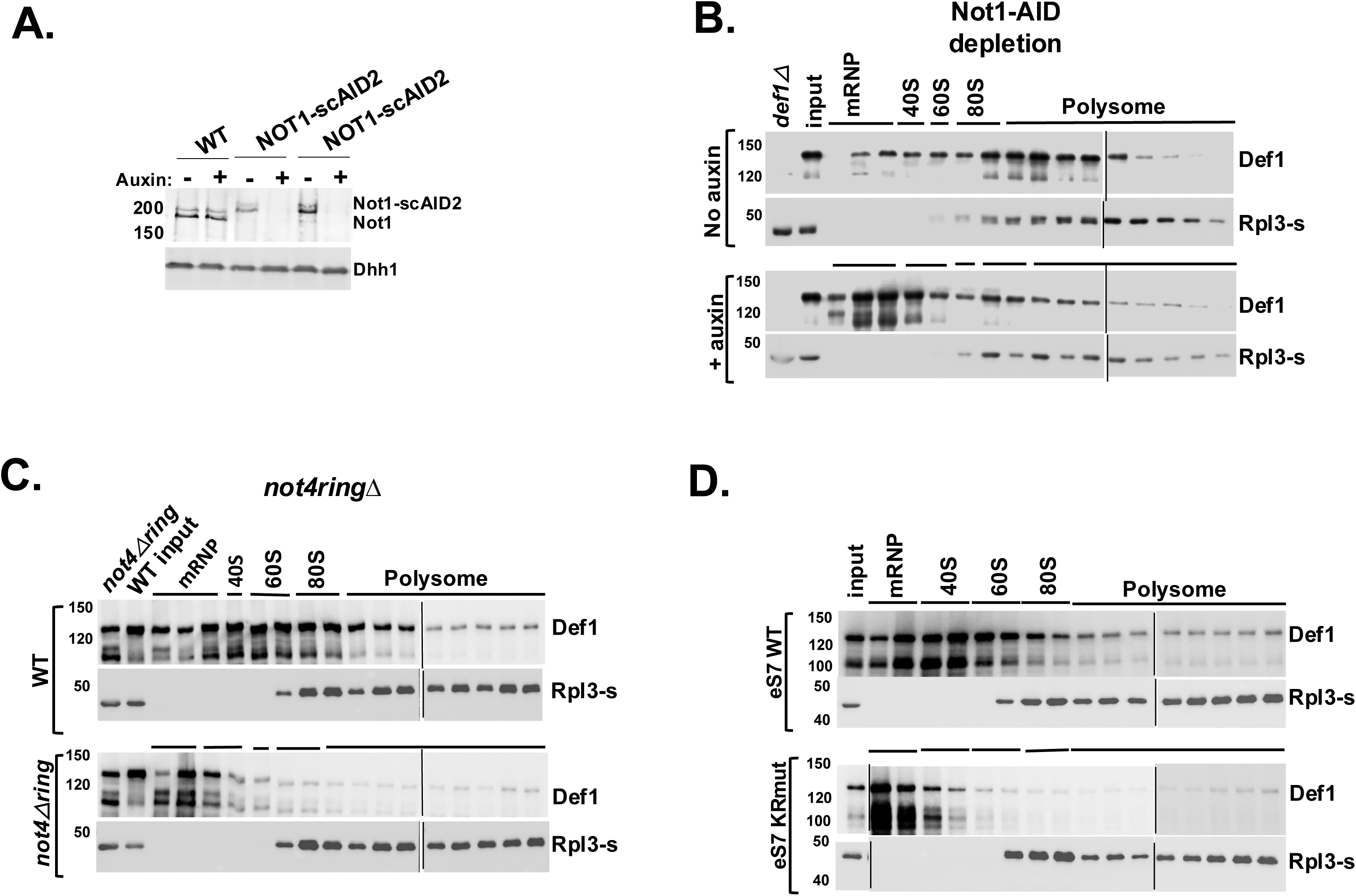
Ubiquitylation of eS7a by Not4 recruits Def1 to ribosomes. (A). Western blot showing Not1 depletion using the auxin-induced degron system. Antibodies to Not1 and Dhh1 (loading control) were used. Two isolates of the Not1-AID strain are shown. Cells were treated with auxin for 2 hr, where indicated. (B). Lysates of untreated and auxin-treated Not1-AID cells were subjected to polysome analysis. Blots were probed with anti-Def1 and Rpl3-s antibodies. (C). Analysis of Def1 binding to polysomes in a *not4Δring* mutant. Blots were probed with anti-Def1 and Rpl3-s antibodies. (D). Lysates from strains expressing wild-type eS7a or a version with lysine to arginine substitution in the Not4-dependent ubiquitylation sites (eS7*_KR_* mut) (Buschauer et al. 2020) were fractionated on sucrose gradients.

Although these results indicate that Ccr4-Not recruits Def1 to ribosomes via Not4’s ubiquitin ligase activity, they only suggest that this is mediated by eS7A (Rps7A) ubiquitylation. These findings might also result from secondary effects arising from disrupting other RING-dependent functions of Not4 or from the modification of other ribosome-associated proteins by Not4. To demonstrate the link between eS7A ubiquitylation and the recruitment of Def1 to polyribosomes, we used a yeast strain that contains lysine-to-arginine substitutions in the ubiquitylation sites modified by Not4 (eS7A-4KR) (Buschauer et al. 2020). In this mutant, we observed a significant loss of Def1 association with ribosomes and an increase in the mRNP fraction, which resembled the *not4RINGΔ* condition (Fig. 6D). This result further supports our conclusion that Not4-mediated ubiquitylation of eS7A is a prerequisite for Def1 recruitment to polyribosomes, highlighting the importance of ubiquitin signaling in this process.

### Def1 recruits the translation and decay factor Dhh1 to ribosomes

Def1, which lacks the C-terminal polyQ domain, associates with polyribosomes (see Fig. 5B). However, this mutant exhibits phenotypes consistent with impaired translation and co-translational decay of mRNAs (Fig. 5D, E, and Supplemental Fig. S3). We hypothesized that the polyQ region recruits proteins to ribosomes, thereby influencing translation or co-translational decay. To test this hypothesis, we performed proximity labeling in strains expressing Def1 variants with increasingly truncated C-termini, each fused to the turbo ID enzyme (TID). The constructs used included Def1_1-380_, Def1_1-500_, and Def1_1-530_, allowing us to identify proteins that associate with the polyQ region of Def1. A growth spot assay demonstrated that fusing TID to full-length Def1 did not slow growth or increase sensitivity to DNA-damaging agents. However, shortening the C-terminus led to phenotypes similar to those of the deletion mutant, as reported by others (Wilson et al. 2013b) (Supplemental Fig. S5A). The expression levels of all Def1-TID fusions were found to be equivalent (Supplemental Fig. S5B). Mass spectrometry identified 97 enriched proteins that were labeled by full-length Def1_1-738_. Of these, 42 proteins were associated with the post-transcriptional regulation of RNA (Fig. 7A and Supplemental File 2). Interestingly, truncating the polyQ region resulted in a significant decrease in the percentage of proteins involved in mRNA regulation (Fig. 7A and Supplementary File 2). The full-length protein labeled 42 RNA regulators, accounting for 43% of the total. In contrast, the truncated versions—Def1_1-380_, Def1_1-500_, and Def1_1-530_—labeled only 15 (11%), 10 (6%), and 11 (9%) RNA regulators, respectively (Fig. 7A and Supplemental File 2).

**Figure 7.**
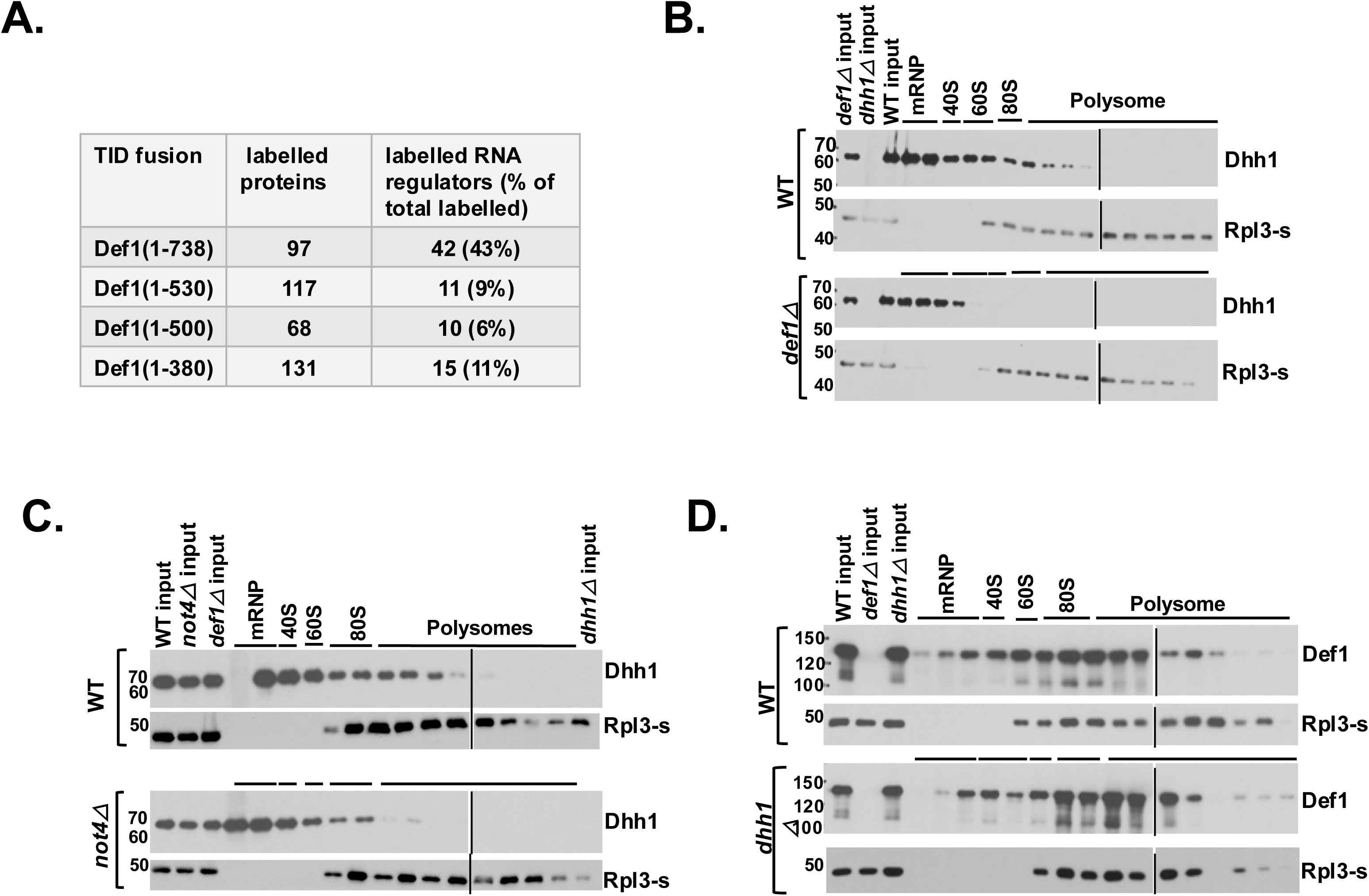
The poly-glutamine-rich (polyQ) domain of Def1 mediates its interaction with mRNA regulators and recruits the translation regulator Dhh1 to ribosomes. (A). BioID was conducted with Def1 truncation mutants fused to TID. Labeled proteins that were enriched for each Def1 derivative versus free TID (>1.5-fold, p<0.05) were tabulated. The number and percent of RNA regulators and ribosome-associated factors identified in the SGD database were calculated and displayed in the table. Dhh1 binding to ribosomes in a *def1Δ* (B) and *not4Δ* (C) mutants. (D). Def1 binding to ribosomes in a *dhh1Δ* mutant. The sample in later polysomes in the *dhh1Δ* mutant was lost during processing in this repetition.

The loss of interaction between RNA regulators and the truncated versions of Def1 suggests that Def1’s C-terminus is responsible for recruiting factors to the ribosome. To explore this further, we analyzed the BioID dataset to identify any interacting proteins that might influence ribosome dynamics or co-translational mRNA degradation. Notably, we found that the DEAD-box RNA helicase Dhh1, a translation repressor involved in co-translational mRNA decay (Coller et al. 2001; Sweet et al. 2012; Radhakrishnan et al. 2016), was labeled by the full-length Def1 protein but not by any of the C-terminal truncated versions. Furthermore, Dhh1 is a good candidate to explore because it plays an important role in codon-dependent co-translational mRNA decay (Radhakrishnan et al. 2016).

We conducted polysome profiling to assess whether the association of Dhh1 with polysomes is dependent on Def1. Our results showed that Dhh1 was associated with polyribosomes and was more enriched in the early polysome fractions, as reported by others (Fig. 7B)(Sweet et al. 2012). Importantly, the migration of Dhh1 with the 80S fraction and polyribosomes was greatly reduced in the *def1Δ* mutant (Fig. 7B). This indicates that Def1 plays a role in recruiting Dhh1 to ribosomes.

Another study showed that Dhh1 binding to ribosomes required Not4 (see the supplement of (Buschauer et al. 2020)). Given that Not4-mediated ubiquitylation is necessary for Def1 to associate with ribosomes, we investigated this further. Our results confirmed that the interaction between Dhh1 and polyribosomes was diminished in the *not4Δ* mutant (Fig. 7C). Next, we explored whether Dhh1 is necessary for Def1 to bind to polysomes. We performed polysome profiling on lysates from the *dhh1Δ* mutant and found that Def1 still sedimented with polysomes in the absence of Dhh1. This observation suggests that Def1 recruitment occurs prior to that of Dhh1 (Fig. 7D). As noted above, tethering Def1-MCP to a reporter mRNA reduced its expression, a function that depended on the C-terminus of Def1 (Supplemental Fig. S3). To further support the hypothesis that Def1 represses mRNA expression, at least partially through Dhh1, we repeated the tethering assay in a *dhh1Δ* mutant background. Our findings indicated that Def1-MCP-mediated repression of the reporter construct required Dhh1 (Supplemental Fig. S6).

### Discussion

An emerging theme in gene regulation is that gene regulatory factors can influence both transcription and mRNA decay in both the nucleus and the cytoplasm.. Yeast proteins Xrn1 and the Ccr4-Not complex are two well-studied examples of this phenomenon (Haimovich et al. 2013; Sun et al. 2013; Chattopadhyay et al. 2022). Our recent research demonstrated that Def1, previously thought to be involved only in DNA repair and the regulation of RNA polymerase II, also has post-transcriptional functions in the cytoplasm, specifically, regulating mRNA half-lives (Akinniyi et al. 2025). The mechanism of how Def1 controlled mRNA turnover remained unknown. Here, we expand on this observation and propose that Def1 is crucial for the co-translational decay of mRNAs enriched with non-optimal codons. Given that a significant portion of mRNA decay occurs co-translationally (Wu and Bazzini 2023), Def1’s ability to recruit decay factors to ribosomes likely contributes to its role in global mRNA decay.

We propose a model in which the stalling of ribosomes leads to Ccr4-Not recruitment, Not4-dependent ubiquitylation of eS7a and Def1 binding to ribosomes through its CUE domain. This interaction allows Def1 to recruit regulators of translation and/or decay via its C-terminal region (Fig. 8). As a proof of principle, we investigated one factor, Dhh1, that binds to Def1 and demonstrated that Def1 is necessary for its recruitment to polysomes. However, it should be noted that Def1-TID labeled other post-transcriptional regulators, some of which required the C-terminus for the interaction (Supplemental file 2), suggesting that Def1’s ability to recruit multiple mRNA regulators to the ribosome may play a crucial role in the posttranscriptional regulation of mRNAs.

**Figure 8.**
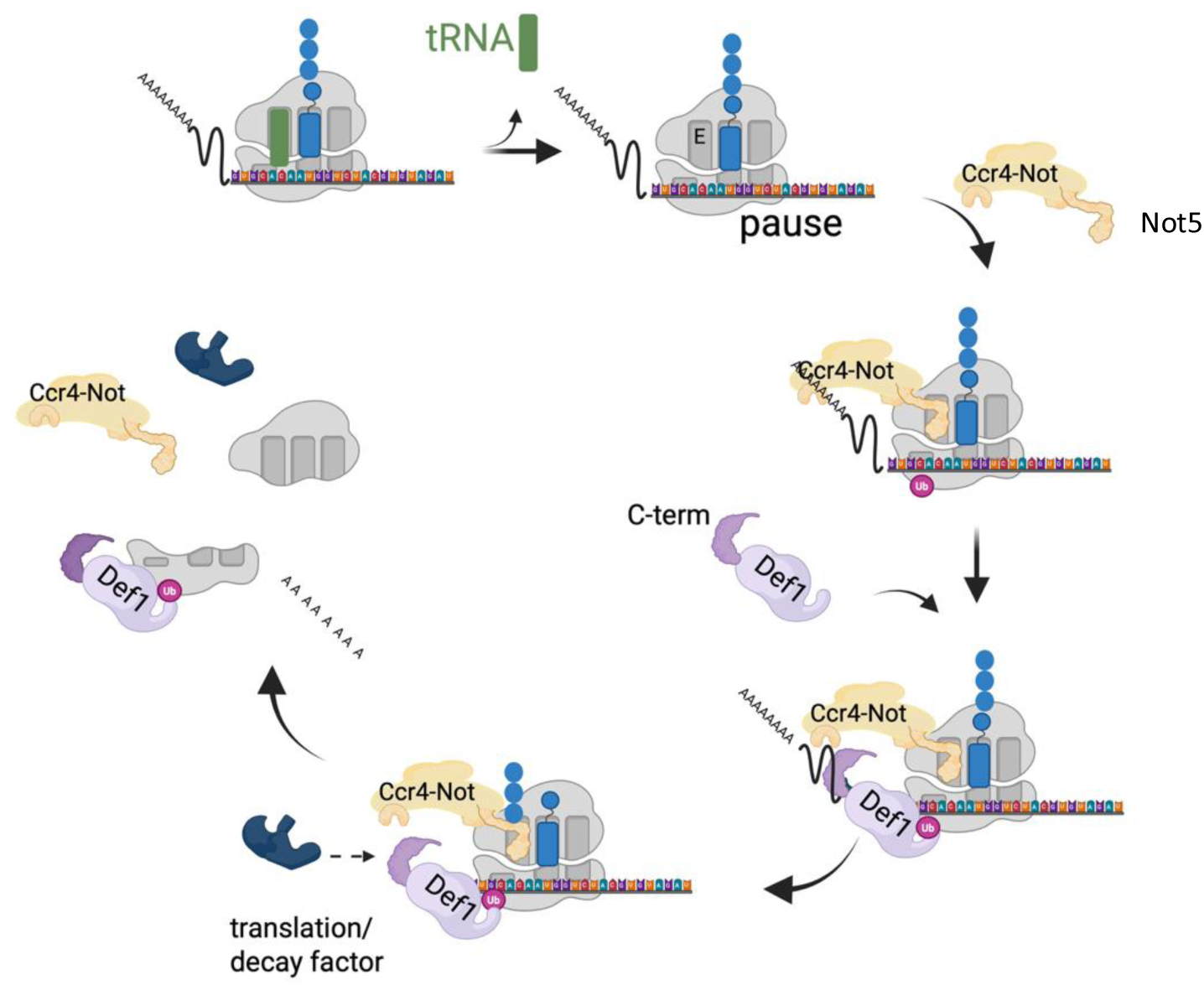
Model for Def1 recruitment to ribosomes. The model is based on the research by Buschauer et al. (Buschauer et al. 2020). When the ribosome experiences arrest or stalling, Not5 binds to the empty E-site, which in turn recruits the Ccr4-Not complex to the ribosome, leading to the ubiquitylation of eS7a. Def1 then binds to the ubiquitylated eS7a through its CUE domain and brings in RNA decay and translation factors to the ribosome via its C-terminal polyQ-rich domain. This recruitment of translation and mRNA regulators facilitates mRNA decay and the removal of the ribosome.

In this context, Def1 acts as a ubiquitin-dependent scaffold that coordinates the recruitment of factors to manage slowly translating ribosomes, which occur due to a high frequency of non-optimal codons. This function resembles its role in degrading Rpb1 during transcriptional stress, where Def1 binds Rpb1 to recruit the elongin-cullin complex and additional factors to the stalled RNA polymerase II (RNAPII) (Wilson et al. 2013b). Recently, the putative human homolog of Def1, UBAP2/2L, was identified for its role in ubiquitylating Rpb1 and in DNA damage resistance (Herlihy et al. 2022).

Interestingly, UBAP2/2L co-sediments with mammalian ribosomes, and knocking it down decreases translation (Luo et al. 2020). Although UBAP2/2L possesses a ubiquitin-binding domain, it lacks the polyQ-rich domain found in Def1, raising questions about whether Def1 has cytoplasmic functions or if UBAP2/2L acquired these functions during the course of evolution. We confirm that Def1 performs both an RNAPII destruction function and a role in translation, like UBAP2/2L. However, there are notable differences between the two proteins beyond the presence or absence of the unstructured polyQ domain. UBAP2/2L is recruited into and is involved in stress granule formation (Youn et al. 2018; Sanders et al. 2020; Riggs et al. 2024), while Def1 does not form detectable foci during stress (Wilson et al. 2013b) and Carillo, R., Akinniyi, O.T. and Reese, J.C, unpublished). Additionally, Luo et al. found that tethering UBAP2/2L to a reporter mRNA increased its expression, whereas tethering Def1 to an mRNA destabilized it and reduced expression (Akinniyi et al. 2025). Here, we also show that Def1 is involved in co-translational decay. Further studies are required to determine whether these differences arise from the specific experimental systems used to study the proteins or from a divergence in their functions.

Def1 influences co-translational decay through a Ccr4-Not-dependent mechanism, but does it have a direct effect on translation? While Def1 is associated with polyribosomes, deleting the gene did not produce any noticeable change in the polysome profiles or the ratio of polysomes to monosomes (Fig. 2C). However, measurements of translation—specifically through puromycin incorporation and changes in the distribution of ribosome footprints across mRNAs—indicate that Def1 does impact translation. The unchanged polysome profile in *def1Δ* cells may result from reduced overall translation rates from initiation to elongation. Ubiquitylation of ribosomal proteins influences multiple stages of translation. Structural studies of the 43S preinitiation complex suggest that ubiquitylation of eS7 disrupts the interaction between eIF4A and eIF4G with the ribosome (Brito Querido et al. 2024). Moreover, deubiquitylation of eS7 is necessary for the cap-binding complex eIF4F to associate with the 43S complex (Takehara et al. 2021). Interestingly, subunits of eIF4A are labeled by Def1-TID (Supplemental file 2). Furthermore, eIF3 straddles both sides of eS7, and cycles of eS7 ubiquitylation and deubiquitylation could dynamically impact eIF3 function by regulating Def1 binding. In this regard, Def1 could sense the translation status of mRNAs from initiation to elongation. Further studies beyond the scope of this work, such as solving the structure of Def1 bound to ribosomes, may provide additional insights.

Ribo-seq showed increased ribosome dwelling over non-optimal codons in the A-site in the null mutant. This observation aligns with the necessity of Not4 ubiquitylation for Def1 binding to polysomes and the role of the Ccr4-Not complex in detecting ribosomes stalled over these codons during elongation (Matsuo et al. 2017; Yan and Zaher 2019; Takehara et al. 2021; Inada and Beckmann 2024). A distinct characteristic of the *def1Δ* mutant, in contrast to *not4Δ* and *not5Δ* mutants, is the reduced ribosome density over approximately 30-40 codons. Interestingly, the number of amino acids translated matches the length of the peptide in the ribosome’s exit tunnel (Wilson et al. 2016). The fate of mRNA is influenced by its nascent polypeptide and associated factors (Höpfler and Hegde 2023). Def1 may perform a function similar to that of the nascent polypeptide complex (NAC). NAC helps delay protein folding in a chaperone-like manner by inserting a domain into the tunnel (Gamerdinger et al. 2019). Notably, the Egd2 subunit of NAC contains a ubiquitin-like domain (UBL), and the CUE domain of Def1 binds UBLs. Additionally, Not4 influences the ubiquitylation and functionality of NAC (Panasenko et al. 2006), further linking the roles of Not4 and Def1 to ribosome function. By binding to ribosomes, Def1 may prevent early termination by the premature recruitment of quality-control factors.

Def1 can be added to the list of factors involved in transcription in the nucleus and translation/decay in the cytoplasm. Ccr4-Not, which we describe as a facilitator for Def1 recruitment to ribosomes, links transcription elongation to translation (Gupta et al. 2016). Def1, too, may be important for nuclear-cytoplasmic communication by coordinating transcription and translation. Def1’s nuclear and cytoplasmic functions may be connected. Agents that cause DNA damage and halt RNA polymerase II also affect RNA and disrupt ribosome activity. The degradation of RNA polymerase II is initiated by transcriptional stress, which can be triggered by depleting GTP pools through the use of 6-azauracil (6-AU) and mycophenolic acid (Wilson et al. 2013b). GTP is vital for ribosome movement, and treatment with 6-AU leads to ribosome accumulation on mRNAs (Gupta et al. 2016). Just as RNA polymerase II detects DNA damage while transcribing a gene, ribosomes can identify untranslatable mRNAs and activate ribosome quality control pathways (Inada and Beckmann 2024). It would be beneficial for cells to detect untranslatable mRNAs in the cytoplasm and halt transcription, as newly synthesized transcripts would face a similar fate. It is plausible that Def1 plays a role in sensing and shutting down gene expression at multiple levels in response to nucleic acid damage.

## Materials and Methods

### Yeast strains and plasmids

Strains and plasmids used in this work are contained in Supplemental Tables 1 and 2, respectively. Gene deletions were produced by homologous recombination using PCR-generated cassettes (Brachmann et al. 1998). CUE domain mutants of *DEF1* were constructed using CRISPR-Cas9 gene editing as described (Laughery et al. 2015). PCR of genomic DNA followed by sequencing of the PCR fragments was used to screen the mutants. Truncation of Def1 was accomplished by inserting GFP at various locations (Longtine et al. 1998). The *HIS3* codon optimality reporters were constructed by replacing the *GAL1* promoter in pJC857 (0% OPT), pJC860 (30% OPT),pJC862 (50% OPT) and pJC867 (100% OPT) with the *GPD1* promoter (Radhakrishnan et al. 2016).

Details about plasmid construction and sequences are available upon request.

### Growth spot assay

Saturated cultures were serially diluted in sterile distilled water and then spotted on plates containing the appropriate media and supplements. Plates were incubated at 30°C for 24 hr or more, as indicated in the respective figures.

### Auxin-inducible degron rapid depletion

Auxin-inducible degron (AID) strains were generated using plasmid constructs as previously used and described (Shetty et al. 2019). Cells were grown to mid-log (OD600∼0.5-0.6) phase in 1% yeast extract, 2% bacto-peptone, 2% dextrose supplemented with 20 ug/mL adenine sulfate (YPAD). Auxin (Indole-3-acetic acid, IAA) was added to a concentration of 500 uM for 2 hr. Protein depletion was monitored by western blotting of proteins isolated by TCA extraction.

### Western blotting and antibodies

Cell extracts were fractionated on either Tris-acetate gradient gels or by SDS-PAGE. The gels were transferred onto a nitrocellulose membrane, and the proteins were visualized by staining with ponceau S. After destaining, the membranes were blocked in 5% dry milk prepared in TBST (50 mM Tris-HCl, pH 7.5, 0.15 M NaCl, 0.1 mM EDTA, and 0.1% Tween-20). Antibodies to Not1 (N-terminus), Not4, Taf14, and Dhh1 are described in other publications (Dutta et al. 2011; Jiang et al. 2019; Pfannenstein et al. 2024). The Def1 antibody was raised in rabbits to a 6HIS-Def1 (1-500) (Akinniyi et al. 2025). Antibodies were validated by probing extracts from null mutants (Dhh1, Not4, and Taf14) or by assessing a shift in migration with a tagged version of the protein (Not1). Examples are displayed in the figures of this paper.

Commercial primary antibodies used are Anti-GFP (UBPbio, Y1031), Anti-FLAG M2 (Millipore Sigma), Rpl3s and anti-puromycin (DHSB hybridoma bank, U. Iowa) and anti-HA, (HA.11, Biolegend). Signals were detected by fluorescence or chemiluminescence (ECL) and imaged on X-ray film or a ChemiDoc-MP imaging system (Bio-Rad).

### BioID experiments

Turbo ID (TID) (Branon et al. 2018)was fused to the C-terminus of Def1, and truncation derivatives, using PCR-derived cassettes from pJR106 as described in (Pfannenstein et al. 2024). The control used in the BioID studies was cells expressing free TID from the *CHA1* promoter. Protein extraction and isolation on streptavidin magnetic beads (Pierce, Thermo Fisher, #88817) was conducted as described in (Kulkarni et al. 2025). For each strain, biological duplicates were analyzed. Proteins were reduced with DTT, alkylated with iodoacetamide, and then digested on the beads with trypsin (Promega Trypsin Gold) overnight at 37°C. The peptides were purified on C18 columns and labeled with TMTpro 16plex reagents (Thermo Fisher, #A44520) per the manufacturer’s recommended protocol. Details on the conditions and instrumentation for mass spectrometry are described in (Akinniyi et al. 2025; Kulkarni et al. 2025).

### Polysome profiling

Fifty ml of mid-log-phase cultures were treated with 100 ug/mL cycloheximide for 10 min. Cells were harvested by centrifugation at 4000 rpm for 5 min. Lysate preparation was carried out as previously described(Panasenko and Collart 2012).

Briefly, cells were resuspended in 500 uL polysome buffer with cycloheximide (CHX) and protease inhibitors (20 mM HEPES-OH pH 8, 10 mM MgCl_2_, 50 mM KCl, 1% triton X-100, 1 mM DTT, 100 ug/mL CHX, 1 mM PMSF, 0.5 ug/mL leupeptin, 1 ug/mL pepstatin) followed by the addition of 500 uL glass beads. Cells were broken by shaking for 18 min on a vortex mixer at 4°C (3 cycles of 6 min bead-beating and 5 min of cooling on ice). Lysates were centrifuged for 10 min at 14,000 rpm and the protein concentration of the clarified supernatant was quantified by Bradford assay. Extracts were stored at -80°C until use. Continuous 10-50% sucrose density gradients (SDG) were prepared on GradientMaster (Biocomp) and stored at 4°C for at least 1hr. 1 mg of protein equivalent of lysate was layered onto the SGD and centrifuged at 39k rpm for 2.5 hr in a SW40 rotor (Beckman Coulter). SDG was fractionated on a Brandel fractionator and the OD254 of peaks were recorded by *PeakChart 3*. Samples used for Western blotting were prepared by precipitation with 10% trichloroacetic acid (TCA) in the presence of 10 or 20 ug of BSA overnight at 4°C. The precipitated proteins were collected by centrifugation at 14k rpm for 10 min and the pellet was washed with 100% acetone. Proteins were solubilized in 1X SDS-PAGE loading buffer and boiled for 5 min. Western blotting was performed as described above. Freezing of the samples was avoided, as this led to Def1 aggregation and uneven signals on the blots.

### Riboseq and data analysis

Ribo-seq was conducted as described previously (McGlincy and Ingolia 2017; Vijjamarri et al. 2023) in triplicate (biological). Cells were not treated with cycloheximide, but it was contained in the extract buffer. After library preparation, 50bp paired-end reads were sequenced. Using fastp, we trimmed the Illumina universal adapter sequence and low-quality bases from the data. Reads with a minimum length of 20bp were retained. Samples were split into replicates based on barcode sequences. Data was mapped to the yeast genome using hisat2 and the RPF read counts at coding sequence (CDS) of each gene was obtained with featureCounts. The CDS raw RPF read counts were used for differential expression analysis. We studied the ribosome profiling features using RiboMiner v0.2 (Li et al. 2020). Transcript sequences were prepared using the prepare transcript command in RiboMiner, and Read1 data were mapped to the transcripts using bowtie2. Frame distribution, length distribution, and periodicity were checked. Based on the periodicity plots, only 28 bp long reads were used for further analysis. P-site was estimated to be at 12 bp offset from the 5 bp end of 28 bp long reads. Metagene analysis was performed, and plots were generated for the entire transcript region and a 500 bp region around the CDS start and end. Polarity values were computed to judge whether the ribosome density is enriched at the 5’ end or the 3’ end. Additionally, codon density at A-sites and P-sites was calculated and plotted. Deseq2 was used to model the RPF and RNA-seq read counts in a generalized linear model (GLM). Genes with significant changes in translational efficiency were obtained using DESeq2 following the DeltaTE approach.

### mRNA decay reporter assays

The codon optimality reporter assays were conducted by transforming yeast with *GPD1-HIS3* plasmids of 0-, 30-, 50- and 100% optimality (see Supplementary Table 2). Cells were grown in -uracil dropout media containing 2% dextrose to an OD_600_ of 0.6-0.8. RNA was isolated by acid-phenol extraction. RNA was subjected to Northern blotting using random-primed biotinylated DNA probes (BioPrime Kit,Thermo-Fisher) to the *HIS3* ORF. Since each the sequence of each *HIS3* open reading frame was different, probes were made for each *HIS3* version used in Northern blotting. RNA was detected using streptavidin-IR800 (Li-Cor) and scanned on a Chemidoc-MP (Bio-Rad). Blots were stripped of the *HIS3* probe and then probed with the *GPD1 ORF* control. The *HIS3* reporter signal was normalized to *GPD1*, and the mutant signals were divided by the wild-type signal. The MS2-MCP mRNA tethering assay was conducted as described in previous publications (Sweet et al. 2012). Plasmids expressing MCP fusions with Def1, or fragments of the Def1 coding sequence, were constructed from gene synthesis and PCR-generated fragments using inPhusion assembly (Takara Bio) (Supplemental Table 2). MCP fusion protein expression plasmids were co-transformed with the reporter vector pJC428 into W303 cells. Cells were grown in triplicate in -uracil/-leucine dropout media containing 2% galactose to an OD_600_ of 0.8-1.0. Expression levels of the MCP fusion proteins were verified using anti-FLAG antibodies, and the levels of reporter GFP protein using anti-GFP antibodies. To determine GFP protein levels, a standard curve of an extract was included on the blots. Loading was normalized to Taf14 or TBP levels, which served as a loading control.

## Data availability

Genomics data has been deposited to GEO under accession number GSE290861). Proteomics data were uploaded to MassIVE (MSV000097607).

## Acknowledgements

Dr. Tatiana Laremore performed Mass Spectrometry at the Penn State Huck Proteomics and Mass Spectrometry Core Facility (RRID:SCR_024462). The Genomics Research Incubator (RRID:SCR_024530) and Genomics (RRID:SCR_023645) cores of the Huck Institutes of Life Science were used in this work. We thank Fred Winston, Jeff Coller, and Toshifumi Inada for providing plasmids and strains. Rachael Green is acknowledged for input on metagene plots of RPFs. This research was supported by funds from National Institutes of Health (R35 GM136353 to J.C.R).

## Author contribution Statement

O.T.A. and J.C.R. conducted wet bench experiments. O.T.A., A.S., S.K., and I.A. provided bioinformatics support and analysis. O.T.A. prepared the figures. O.T.A. and J.C.R. wrote the paper.

