## supplemental figures for "Degradation factor 1, Def1, regulates mRNA translation and decay through Ccr4-Not-dependent ubiquitylation of the ribosome"

Supplemental Figure S1

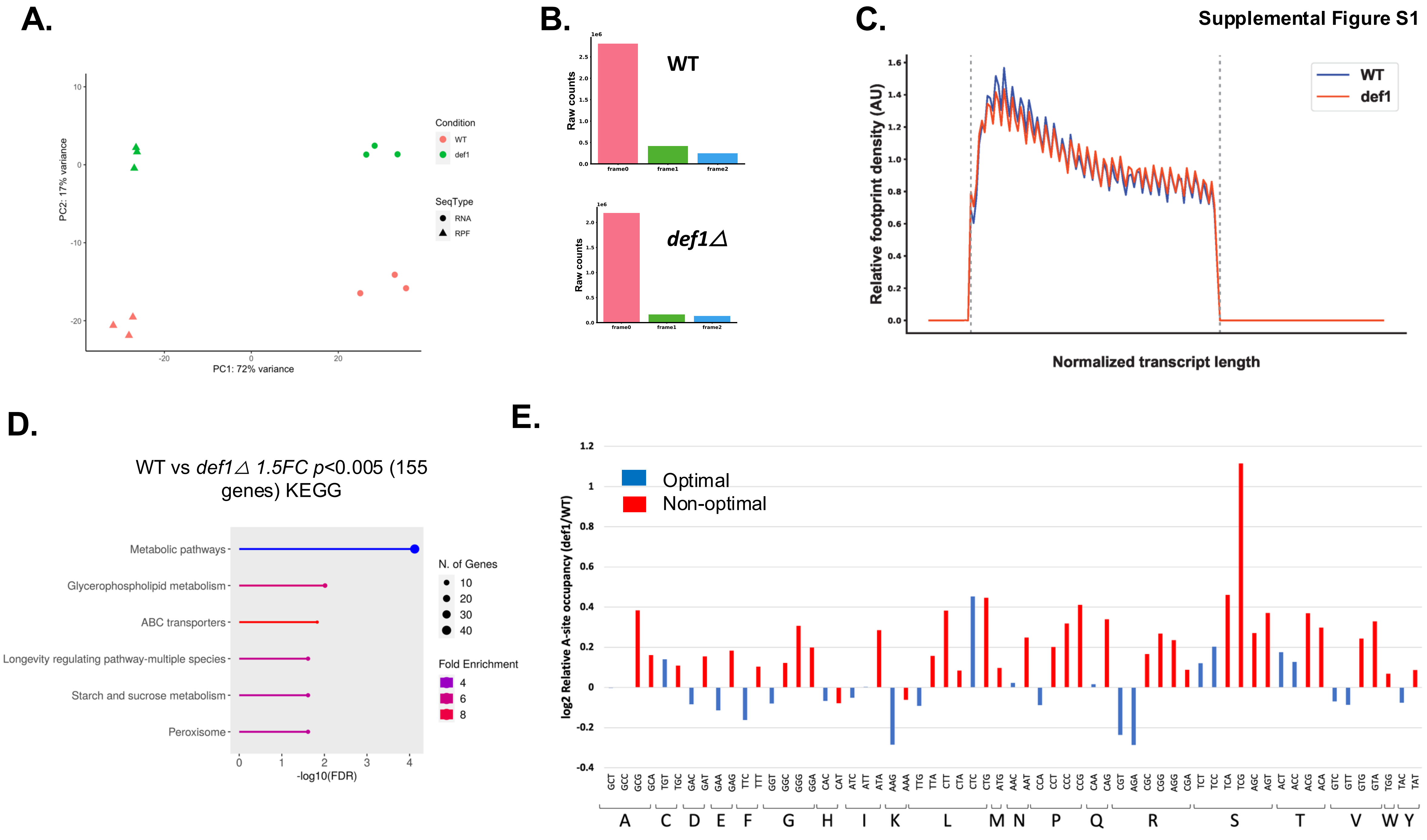

Supplemental Figure S2

A.

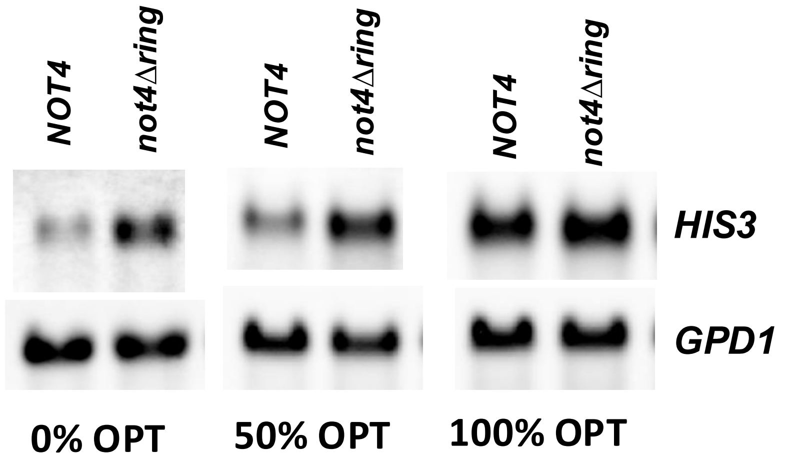


B.

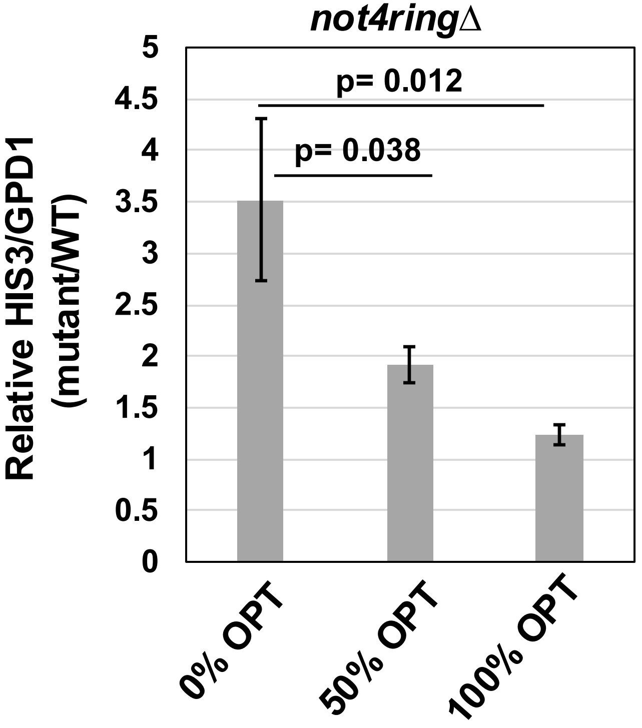

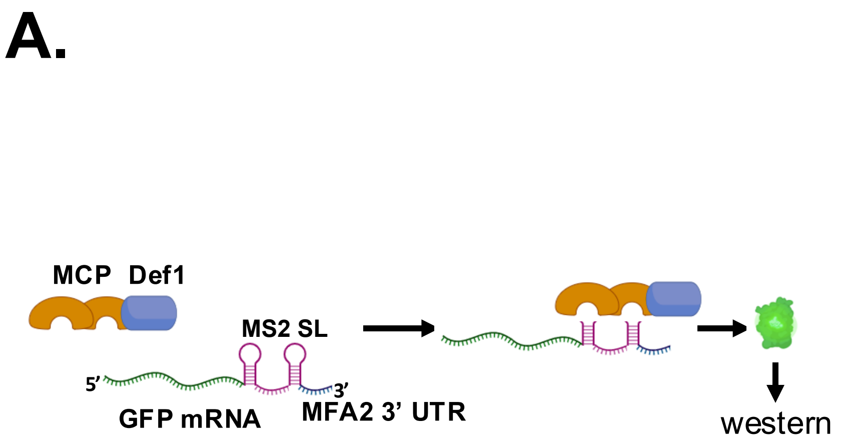
A.
MCP Def1
MS2 SL
5'
3'
GFP mRNA
MFA2 3' UTR
western

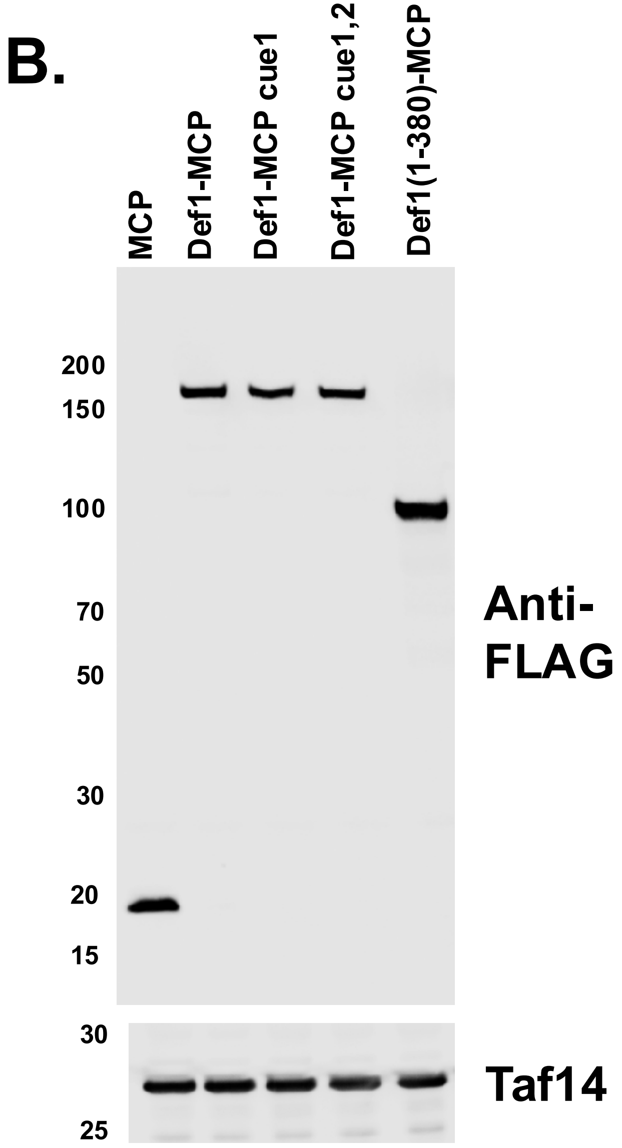
B.
MCP
Def1-MCP
Def1-MCP cue1
Def1-MCP cue1,2
Def1(1-380)-MCP
200
150
100
70
50
30
20
15
Anti-FLAG
30
25
Taf14

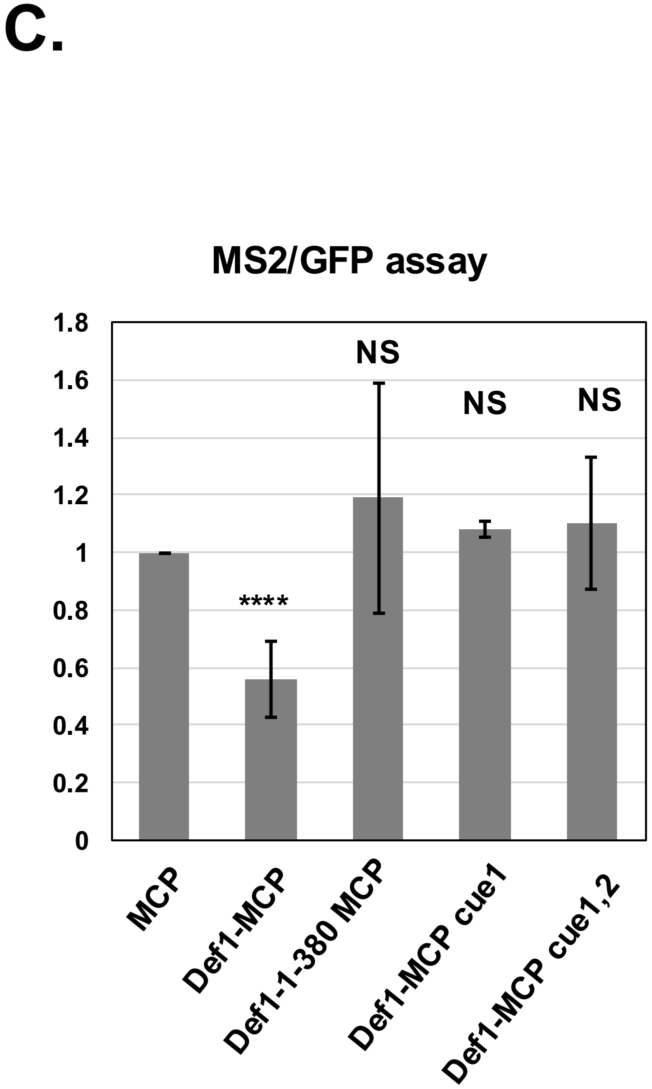
C.
MS2/GFP assay
1.8
1.6
1.4
1.2
1
0.8
0.6
0.4
0.2
0
NS
NS
NS
****
MCP
Def1-MCP
Def1-1-380 MCP
Def1-MCP cue1
Def1-MCP cue1,2

Supplemental Figure 4

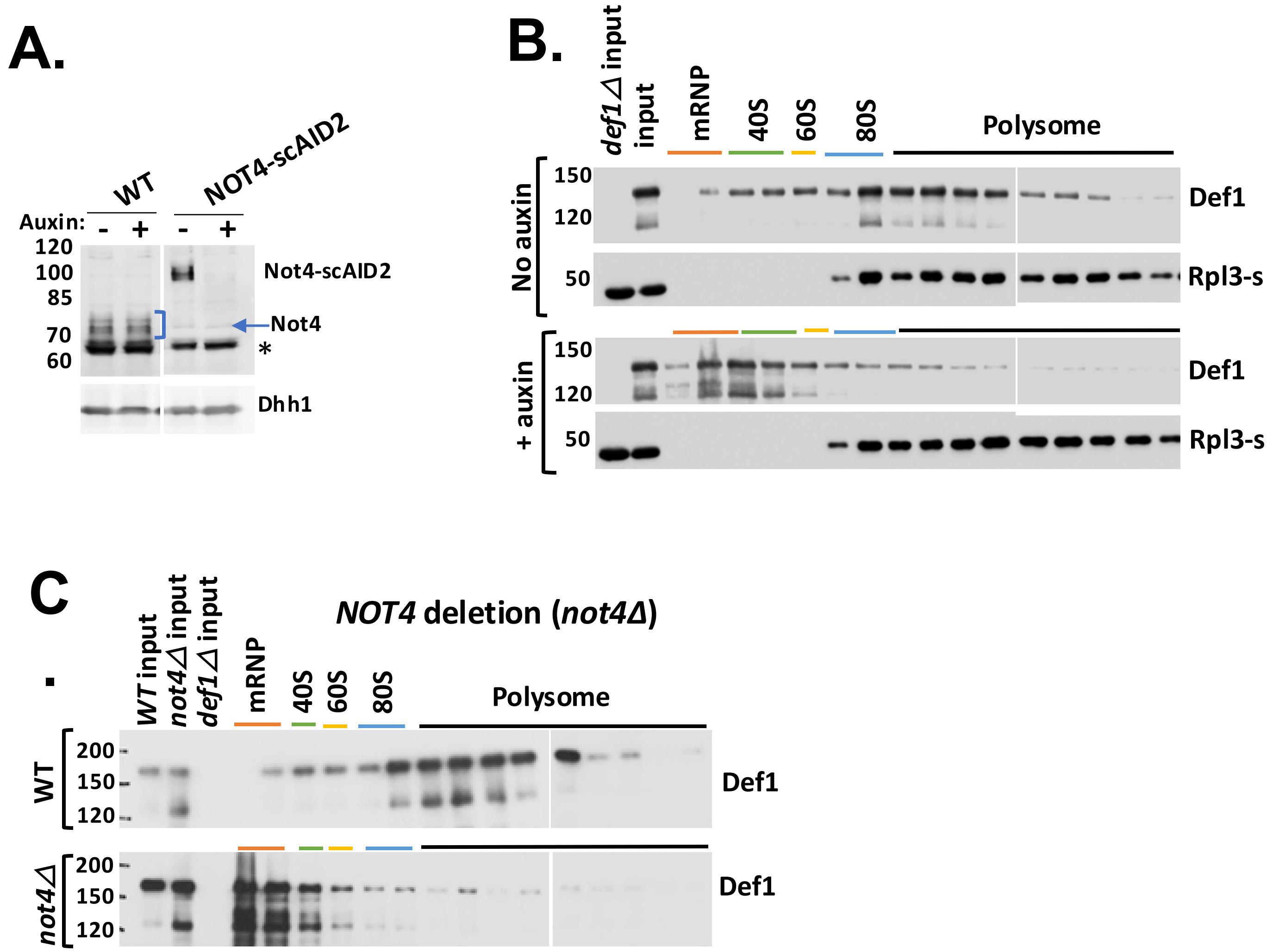

Supplemental Figure 5

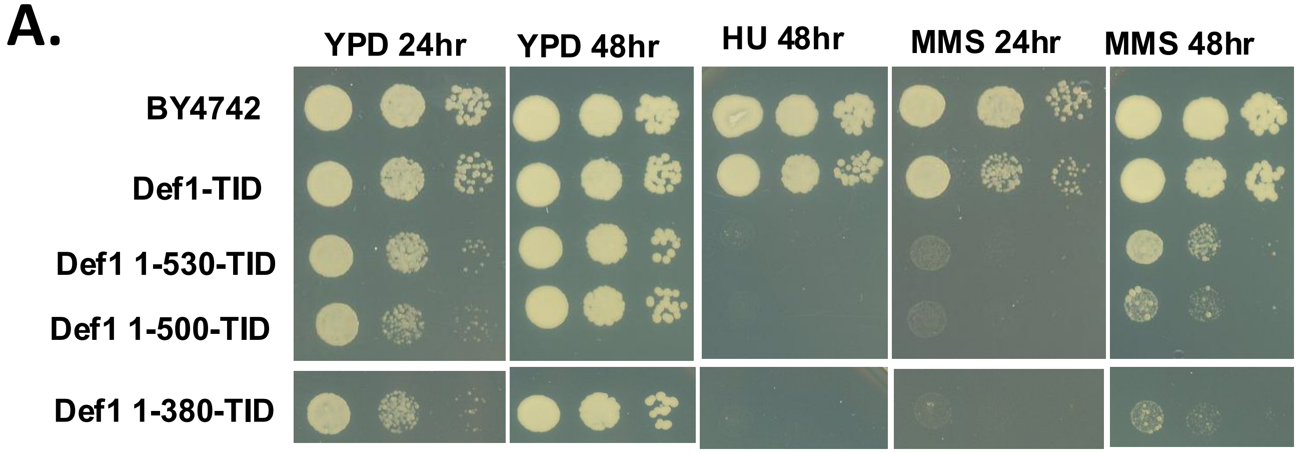


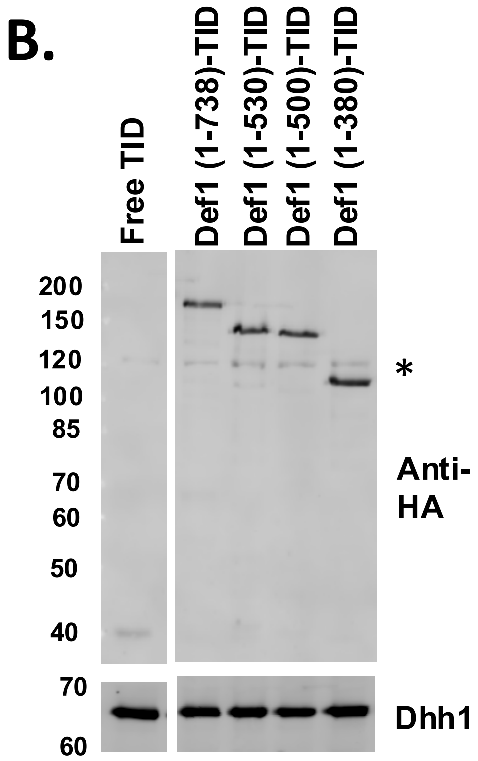

Supplemental Figure 6

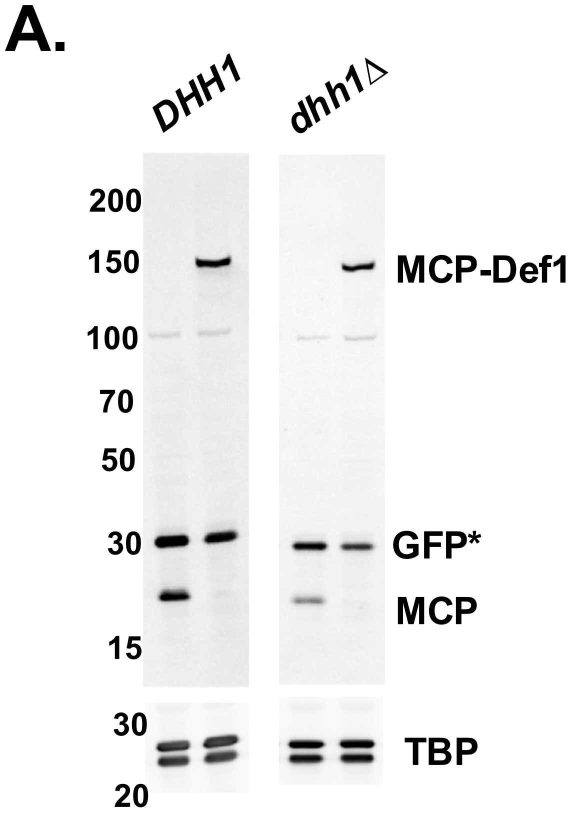


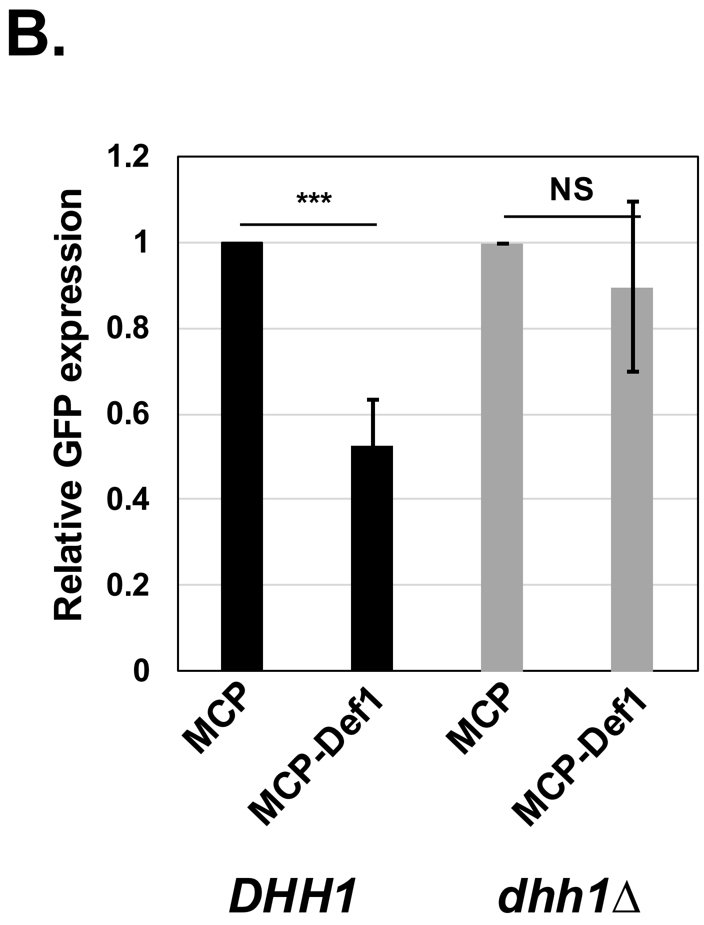

**Supplemental figure legends**

**Supplemental Figure 1**: Ribo-seq supplemental information. (A). Principal component analysis displaying ribosome footprint and total RNA data. The RPF and total RNA reads from all 3 biological replicates of each condition were analyzed. (B). Distribution of RPF reads in all three reading frames. RPFs were mapped to reading frames in wild-type and *def1Δ* cells. The relative abundance of in-frame (with respect to the start codon) (Frame0) and out-of-frame (Frame1 and Frame2) was plotted. (C) Metaplots of RPFs. Plots of ribosome protection across transcript ORFs were generated using RiboMiner and normalized to transcript length for wild-type (blue) and *def1*Δ (red) cells. (Li *et al,* 2020) (D). KEGG analysis of the 155 mRNAs with significantly increased (>1.5FC p<0.005) translation efficiency (TE). (E). Relative ribosome A-site occupancy of all 61 amino acid codons. Differential occupancy in the A-site (log2) in *def1*Δ versus wild-type cells was plotted. Codons were binned into optimal (blue, CSC>0.47) or non-optimal (red, CSC <0.47) based on CSC scores (Presnyak *et al*., 2015).

**Supplemental Figure 2. Validation of the GPD1 codon-optimality decay assay.**
(A). Wild-type and *not4ring*Δ cells were transformed with plasmids expressing *HIS3* ORFs with variable codon optimality (indicated below each panel) under the control of the *GPD1* promoter. The endogenous *GPD1* mRNA was used as a loading control and to correct for changes in promoter activity in the mutant. The image capture times for each *HIS3* derivative varied because of differences in steady-state levels. (B). Averages and standard deviations of reporter mRNA expression. The signals for the *HIS3* mRNA were first normalized to the endogenous *GPD1* mRNA, and then the values in the *not4ring*Δ cells were divided by the values in wild-type cells (relative normalized signal) (n=3, biological triplicates). P-values were calculated using a two-tailed, unpaired t-test, and the values of the 0% reporter were compared to those of the other constructs.

**Supplemental Figure 3. MS2/MCP mRNA tethering assay.**
(A). Graphic of the MS2/MCP assay. (B). Expression of MCP fusion proteins, blotting with anti-FLAG M2 monoclonal antibody. Taf14 was the loading control. (C). MS2-MCP tethering assay for GFP protein expression in the Def1-MCP derivatives is indicated in

the panel. The amount of GFP protein in cells expressing “free” MCP was set to 1.0. The averages and standard deviations are displayed. Def1-MCP and Def1(1-380)-MCP samples (N=6) and CUE mutants (N=3) are displayed. ***, p<0.001

**Supplemental Figure 4**: **NOT4 is required for Def1 binding to ribosomes.** (A). Auxin-inducible degron (AID) was used to deplete Not4. Cells were treated with auxin for 2 hr, where indicated. Lysates were subjected to western blotting using anti-Not4 and anti-Dhh1 serum (control). The asterisk marks a protein that cross-reacts with the anti-Not4 antibody(Jiang et al. 2019; Pfannenstein J 2024). Not4 runs as a broad multimer in cells in some gel systems. (B). Fractionation of cell lysates on sucrose gradients from NOT4-AID cells with or without auxin treatment. Fractions were analyzed by western blotting using anti-Def1 antibody. An extract of *def1*Δ is included to validate the antibody. Rpl3-s is a large subunit ribosomal protein. (C). Fractionation of cell lysates on sucrose gradients from *not4*Δ cells.

**Supplemental Figure 5.** (A) growth phenotypes of the Def1-TID strains. Saturated cultures were serially diluted and spotted onto the media indicated in the panel. Fusing TID to full-length Def1 does not cause growth phenotypes. Deleting the C-terminus results in growth defects, as previously described in another publication (Wilson et al. 2013). (B). Expression levels of TID-labelled Def1 truncation mutants by western blotting. Anti-HA antibodies were used. A HA-tag is incorporated into the fusion proteins(Pfannenstein J 2024). Asterisks mark a cross-reacting band. Dhh1 serves as a loading control.

**Supplemental Figure 6. MS2/MCP reporter assay in *dhh1*Δ cells.** (A). Western blotting using anti-FLAG (M2) antibodies to measure fusion protein expression. TBP is the loading control. The GFP signal is due to incomplete stripping of the blot before reprobing with anti-FLAG antibodies. (B). GFP expression. Assay was performed as described in the legend of Supplemental Figure 3.
